# SARS-CoV-2 neutralization and serology testing of COVID-19 convalescent plasma from donors with non-severe disease

**DOI:** 10.1101/2020.08.07.242271

**Authors:** Thomas J. Gniadek, Joshua M. Thiede, William E. Matchett, Abigail R. Gress, Kathryn A. Pape, Marc K. Jenkins, Vineet D. Menachery, Ryan A. Langlois, Tyler D. Bold

## Abstract

We determined the antigen binding activity of convalescent plasma units from 47 individuals with a history of non-severe COVID-19 using three clinical diagnostic serology assays (Beckman, DiaSorin, and Roche) with different SARS-CoV-2 targets. We compared these results with functional neutralization activity using a fluorescent reporter strain of SARS-CoV-2 in a microwell assay. This revealed positive correlations of varying strength (Spearman r = 0.37-0.52) between binding and neutralization. Donors age 48-75 had the highest neutralization activity. Units in the highest tertile of binding activity for each assay were enriched (75-82%) for those with the highest levels of neutralization.

## Background

The transfer of passive immunity with convalescent plasma is a promising strategy for treatment and prevention of COVID-19 [1]. Challenges include the heterogeneity of anti-SARS-CoV-2 antibody responses [2] and the variety of assay techniques to measure antigen binding activities [3]. Although greater severity COVID-19 disease is correlated with antibody titers [4], the majority of individuals diagnosed with COVID-19 (and potential convalescent plasma donors) have non-severe disease [5]. It is therefore especially important to assess the variability in antigen binding activity of convalescent plasma from donors with less severe disease and to determine how readily available serologic measurements correlate with functional neutralization activity in this population.

Viral neutralization titers measure the ability of antibodies to prevent viral infection of a eukaryotic cell line in vitro. Both live and pseudotyped virus assays exist for SARS-CoV-2 [6,7], as well as surrogate assays that measure blockade of the Spike-ACE2 protein interaction [8]. Pseudotyped and recombinant protein assays require less restrictive biosafety procedures and facilities; however, results may differ from live viral assays since pseudotyped viruses typically express only a single viral entry protein. Neutralizing antibodies that impact other viral targets and processes may therefore not be detected by pseudotyped viral assays [9]. Readouts of neutralization assays also vary, but standard viral plaque reduction readouts are labor-intensive. Fluorescence-based assays use an engineered viral particle with a fluorescent protein gene expressed upon viral infection of eukaryotic cells [10]. Fluorescence assays correlate well with plaque reduction, but are faster and readily automated[7]. Despite these innovations, live SARS-CoV-2 neutralization assays remain available only in specialized laboratories and are not widely used to screen convalescent plasma due to complexity of implementation, high cost, low throughput, and biosafety concerns.

In addition, clinical assays that measure the binding of antibodies against SARS-CoV-2 to specific viral antigens have been rapidly developed. These assays were primarily designed to diagnose past COVID-19 exposure and none of the currently available assays measure the ability of antibodies to neutralize the virus or prevent viral entry into cells. However, many of these clinical assays can be run on high-throughput, automated instruments. As a result, these assays can be performed at low cost, low safety risk to laboratory staff, high throughput, and in almost any clinical laboratory.

Initial reports have suggested some correlation between binding antibody activity and viral neutralization titers [11]. Data from other coronaviruses suggests that plasma with detectable antibodies at a 1:160 or 1:320 dilution using a clinical binding assay should have high neutralizing titers as well [11]. However, these assumptions have not been rigorously tested and the dilution of samples requires specific assay validation and overhead. Even without dilution, many clinical serology assays provide a quantitative serologic score of antibody reactivity.

Despite these uncertainties, there is currently high demand for COVID-19 convalescent plasma for compassionate use and clinical trials. Therefore, we sought to determine in a real-world cohort of donors with a history of non-severe COVID-19, the relationship between antigen binding activity measured by several FDA approved clinical diagnostic assays, and neutralization activity against live SARS-CoV-2 recombinantly engineered to express fluorescent mNeonGreen protein in infected Vero E6 cells [7,10].

## Methods

### Convalescent Plasma Donor Recruitment

The NorthShore University HealthSystem COVID-19 convalescent plasma collection program was established in April 2020. This program and associated human subject research, performed in in accordance with the ethical standards of the Helsinki Declaration, were approved by the NorthShore University HealthSystem Institutional Review Board. All potential donors provided written consent for the study and provided information about their COVID-19 disease history and demographics. Disease history was reported in free-text format and the absence of a reported symptom was assumed to indicate that the symptom was not present. For the donors included in this study, reported symptoms included fatigue (49%), myalgia (47%), cough (47%), anosmia (43%), headache (40%), and other symptoms <20% each (Table S1). None of the donors had a history of hospitalization for COVID-19.

### Sample Collection, Storage, and Transport

Samples were collected at the time of donation using serum separator tubes (BD, Franklin Lakes, NJ), centrifuged, aliquoted, and frozen at −80 C. For each sample included in this study, an aliquot was shipped to the University of Minnesota for viral neutralization titer measurement on dry ice.

### Clinical Serology Testing

An aliquot was thawed and tested at NorthShore using the Elecsys Anti-SARS-CoV-2 (Roche Diagnostics), Access SARS-CoV-2 IgG (Beckman Coulter), and LIAISON SARS-CoV-2 S1/S2 IgG (DiaSorin). Both the quantitative cutoff index and qualitative results were recorded. Samples were divided into tertiles based on the Roche assay results, then 47 were randomly selected to equally sample each tertile.

### SARS-CoV-2 S1 RBD Ig ELISA

The anti-SARS-CoV-2 S1 RBD total Ig assay employed a standard indirect enzyme immunoassay technique (ELISA), described in detail elsewhere, using a secondary antibody recognizing all human immunoglobulin isotypes (goat anti-human IgG H+L-HRP, Invitrogen/ThermoFisher) [12,13]. Three-fold serum dilutions were tested: 1:50 to >1:12,150. ELISA titer was reported as the dilution at which absorbance of each sample tested exceed the 50% maximal absorbance signal for a positive control sample on the same ELISA plate.

### Live SARS-CoV-2 Virus Neutralization Assay

Vero E6 cells (2.5×10^4^) were seeded in each well of a 96-well Black/Clear Flat Bottom TC-treated plate (Falcon) and incubated in DMEM overnight at 37°C with 5% CO2 prior to infection. Plasma samples were two-fold serially diluted (from 1:20 to 1:5120) in DMEM and incubated with mNeonGreen SARS-CoV-2 at 37°C for 1 hr. Medium was removed from cells and the virus-plasma mixture was added to achieve a final multiplicity of infection (MOI) of 0.1 plaque forming units per cell. The cells were incubated at 37°C with the virus-plasma mixture for 24-26 hr. Following incubation, cells were fixed in 4% paraformaldehyde (PFA) at 4°C for 30 minutes. The PFA-virus-plasma mixture was removed, cells were washed once with PBS, and 50 μl of PBS was added to each well. The fluorescence signal was determined by reading the plates on a BioTek Synergy H1 Hybrid Multi-Mode Reader, using excitation/emission wavelengths of 488/517 nm. Percent maximal infection was determined for each dilution as the ratio of the fluorescent signal to the maximal signal for non-serum-treated controls in the same plate. A nonlinear regression method was used to determine the dilution that neutralized 50% of mNeonGreen fluorescence (NT_50_) by using Prism 8 (GraphPad). If a plasma titration failed to generate 50% inhibition within the range of concentrations tested, a titer value of ½ (1:10) of the highest serum concentration tested was ascribed to it. Each sample was tested in duplicate.

### Statistical Analysis

Assay results were compared using linear regression and Spearman correlation. Results from each assay were broken into equal tertiles for comparison; tertiles were compared using the Kruskal-Wallis test. Comparisons of NT_50_ between two groups used the Mann-Whitney U test.

## Results

### Fluorescent SARS-CoV-2 neutralization

We tested 47 units of convalescent plasma units from individuals with a history of non-severe COVID-19 for viral neutralization activity. There was a broad range of neutralization against live SARS-CoV-2, with 6 (13%) units demonstrating high NT_50_ values >500, and 6 (13%) with undetectable high NT_50_ values < 20 (Figure S1). Neutralization assays results yielded a robust calculation of NT_50_ with R^2^ values > 0.9 for curve fitting (Figure 1A) in the majority of samples with detectable neutralization activity.

**Figure 1:**
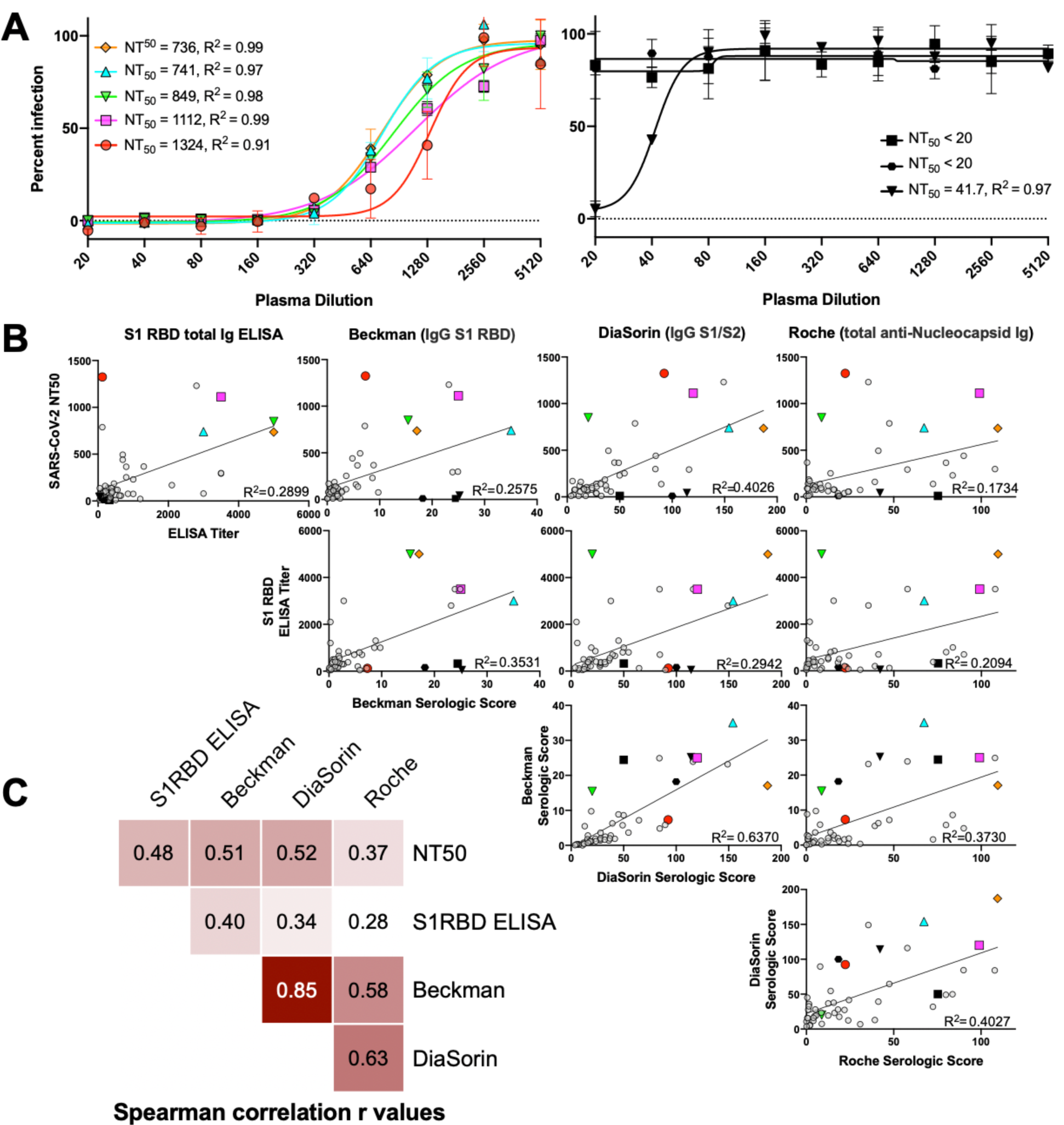
Comparison of non-severe COVID-19 convalescent plasma SARS-CoV-2 neutralization activity and antigen binding assays. (A) Dilution curves with example of non-linear curve fitting showing R^2^ and NT_50_ calculation for five highly neutralizing units and three units with low or undetectable neutralization activity. (B) Correlation plots comparing all assays tested, with linear curve fitting R^2^ value shown. Highly neutralizing samples corresponding to example dilution curves in 1A are represented in all plots with colored symbols, while samples with low or undetectable neutralization are denoted with black, filled symbols. (C) Spearman’s correlation r values for pairwise comparisons of each assay tested.

### Binding and Neutralization Assay Comparison

We compared the neutralization activity in these samples to binding activity as measured by an in-house ELISA for S1 RBD total Ig, as well as three clinical diagnostic assays that use different viral antigenic targets: Beckman (IgG anti-S1 RBD) DiaSorin (IgG anti-S1/S2 protein) and Roche (total anti-Nucleocapsid Ig). There was a universally positive relationship between NT_50_ and all four binding assays tested, with Spearman correlation r ranging from 0.37-0.52, and R^2^ values of 0.17 to 0.40 reflecting a weak linear relationship (Figure 1B and C). The strongest positive correlation between assays was the Beckman and DiaSorin assays (Spearman r = 0.85), which measure different aspects of anti-Spike protein binding activity. The Roche total anti-nucleocapsid assay had the lowest overall correlation with NT_50_, but stronger positive correlation with the Beckman and DiaSorin assays (Figure 1C).

We identified several individual donors with discordant binding and neutralization activity, including some with high neutralization and low binding activities in individual assays (Figure 1A and B, red circles and green inverted triangles). We also identified individual units characterized by low neutralization despite relatively high binding activity in individual assays (Figure 1A and B, black square and inverted triangle).

### Tertile analysis to enrich for highly neutralizing convalescent plasma units

Because the optimal NT_50_ that corresponds to functional immune protection is not known and depend on the neutralization assay used, we assigned each convalescent plasma unit to a neutralization tertile and sought to determine how different donor and binding characteristics enrich for units found in the top two tertiles of neutralization activity. These included units with NT50 values of either 1:78-180, or >1:180. There was a significant difference in NT_50_ depending on donor age. Donors in the oldest tertile (age 48-75) had the highest enrichment for the top two neutralization tertiles (p=0.04). However, no other donor demographic including sex, fever, symptom duration or reported COVID-19 symptoms correlated with NT_50_ (Figure 2A and S2). We observed no association between neutralization or binding and the time from either symptom onset or symptom end to sample collection (Figure S2).

**Figure 2:**
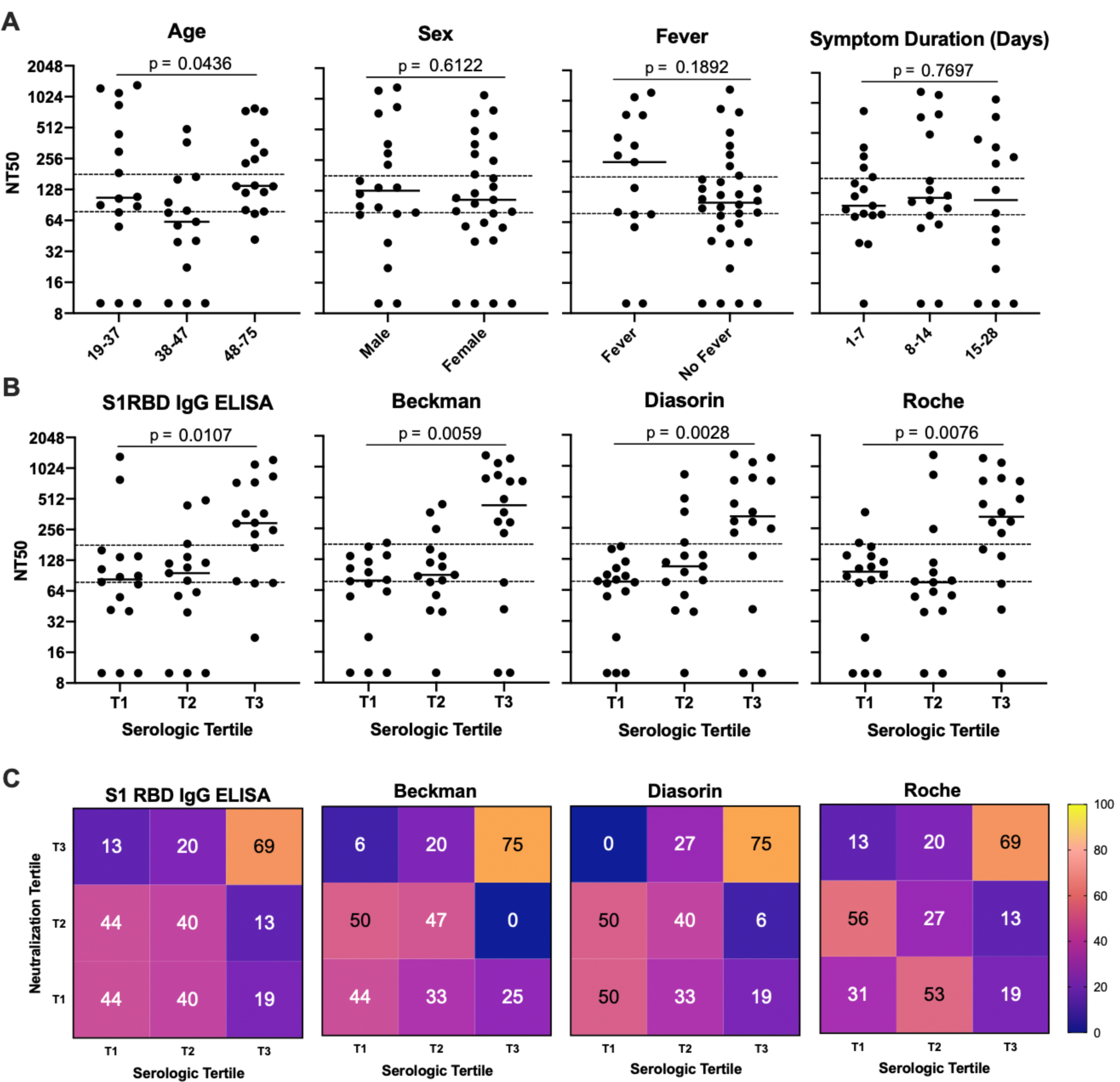
Enrichment for highly neutralizing non-severe COVID-19 convalescent plasma by donor characteristics and serologic tertile. (A) Dot plots demonstrating NT_50_ values for all units tested as categorized by key donor characteristics. Dotted horizontal lines represent transition points between neutralization tertiles. (B) Comparison of NT_50_ values between tertiles of binding activity for each serologic assay tested. (C) Heatmaps indicating the percentage of each binding tertile that is made up of units from each neutralization tertile. Statistical significance was assessed by Mann-Whitney U test for comparisons of two donor categories. Kruskal Wallis test was used to compare groups of three categories.

Most highly neutralizing antibodies were in the top tertile of binding activity for each assay. 69% of the units in the top tertile of binding activity for the S1 RBD ELISA assay were in the highest tertile of neutralization activity and 13% in the middle neutralization tertile (82% were highly neutralizing), see Figure 2B-C. Clinical serological assays were similar in this regard, 75-82% of the units in the highest binding tertile were highly neutralizing. Only the S1 RBD ELISA contained no units with undetectable neutralization activity in the highest binding tertile (Figure 2B); whereas each clinical assay contained at least one non-neutralizing sample in the highest binding tertile.

## Discussion

These findings illustrate the difficulty in deriving information about functional antibody responses using anti-SARS-CoV-2 binding assays. Although a correlation exists, the relatively high discordance rate in donors with a history of non-severe COVID-19 may adversely affect the interpretation of convalescent plasma clinical trial data. Furthermore, the discordance observed implies that SARS-CoV-2 patients develop a broad antibody repertoire against multiple proteins and epitopes; each of which may only partially contribute to the overall neutralization of the virus.

Several larger studies that included hospitalized patients with more severe disease have shown higher anti-COVID-19 antibody binding reactivity in individuals who had been hospitalized or received treatment for COVID-19. Interestingly, anti-COVID-19 antibody levels, as measured by binding assays, are high in hospitalized patients during their hospitalization and formation of these antibodies is not related to a decline in viral load [2]. Future studies are needed to understand the significance of these observations. It is possible that high titer binding antibodies with low neutralization potential lead to antibody dependent enhancement, suggesting that selecting convalescent plasma units based on high binding activity alone may cause harm [14]. Similarly, future studies are needed to determine if developing high titer binding antibodies with low neutralization potential is a poor prognostic sign.

In this study, increasing age correlated with NT_50_, consistent with other reports [15], perhaps due to relatively more severe disease processes and greater viral exposure in this age group, though the same association was not seen with symptom duration or fever. However, given the heterogeneity of COVID-19 disease symptoms, it is unclear whether how well symptom history reflects the degree of immunological stimulation.

Although the neutralizing antibody dose needed for clinical benefit is unknown, units in our top 2 neutralization tertiles had activity consistent with current FDA recommendations (NT50 >1:160 or >1:80). Other factors such as the number of units transfused, the viral load at the time of transfusion, the degree of irreversible end organ damage at the time of transfusion, and the extracellular fluid volume of the recipient may be critical factors in determining whether a unit with a certain concentration of neutralizing antibody shows clinical benefit.

## Footnotes

The authors declare no financial or otherwise beneficial conflicts of interest.

## Acknowledgements

We thank Christine Ronayne for administrative and logistical laboratory support, Dr. Jessica Fiege, PhD, for assistance with cell culture, and members of the UMN BSL3 program.

## Supplemental Materials

**Table S1.** Non-severe COVID-19 convalescent donor characteristics.

| Demographics | Number (Percent) |
| --- | --- |
| Age |  |
| 18-37 | 16 (34) |
| 38-47 | 15 (32) |
| 48-75 | 16 (34) |
| Sex |  |
| Male | 20 (43) |
| Female | 27 (57) |
| PCR diagnosis | 44 (94) |
| Symptoms |  |
| Fever | 15 (32) |
| Headache | 19 (40) |
| Nausea | 6 (13) |
| Fatigue | 23 (49) |
| Chills | 9 (19) |
| Myalgias | 22 (47) |
| Diarrhea | 9 (19) |
| Sore throat | 9 (19) |
| Rhinorrhea | 3 (6) |
| Congestion | 9 (19) |
| Chest tightness | 6 (13) |
| Cough | 22 (47) |
| Anosmia | 20 (43) |
| Night sweats | 4 (9) |
| Dyspnea | 8 (17) |
| Rash | 3 (6) |
| Symptom duration |  |
| 1-7 days | 17 (36) |
| 8-14 days | 16 (34) |
| 15-28 days | 14 (30) |

## Supplemental Figure Legends

**Figure S1.**
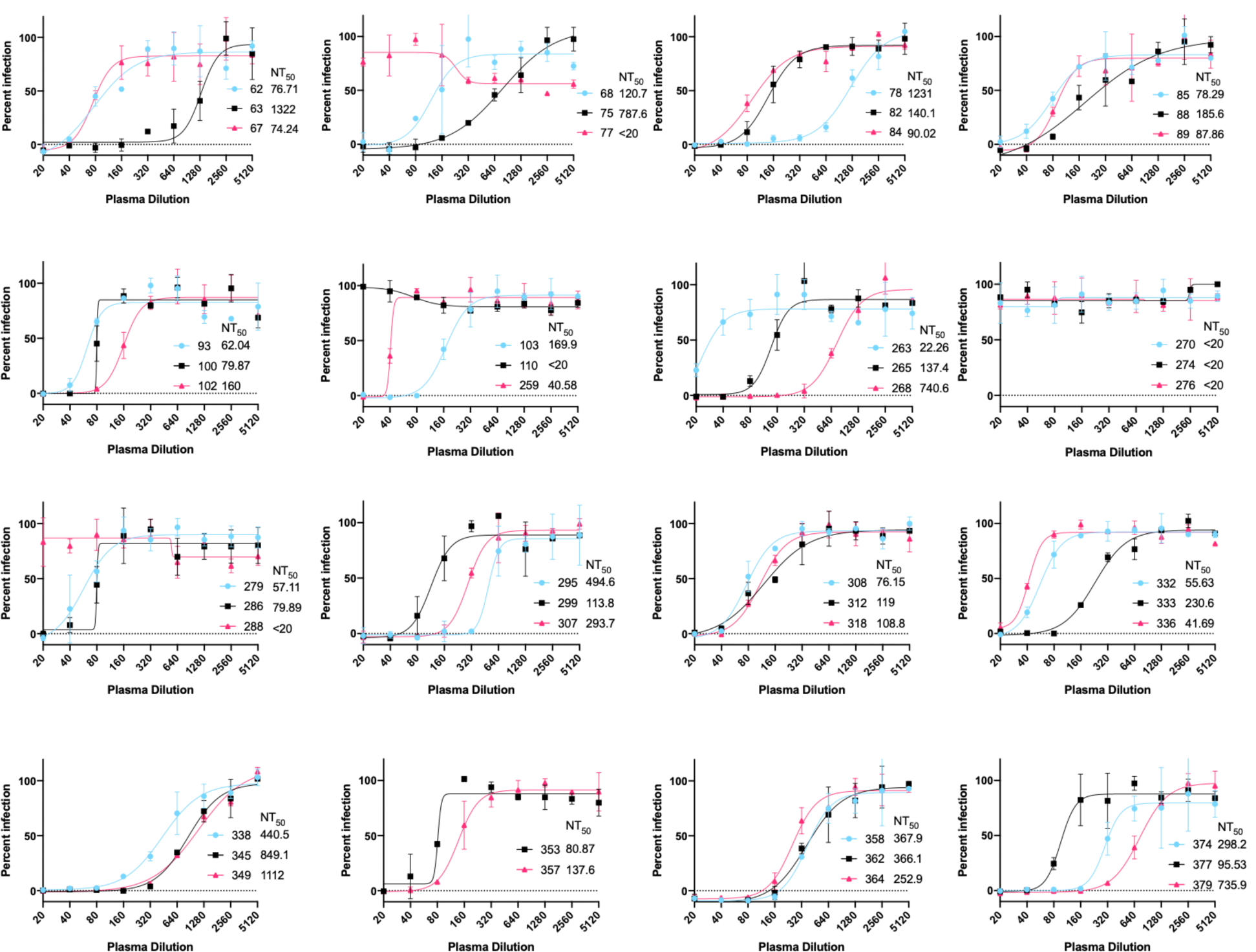
Neutralization curves for all units tested. Dilution series of all 47 plasma units tested are presented in groups of 2-3, with calculated NT_50_ values for each shown.

**Figure S2.**
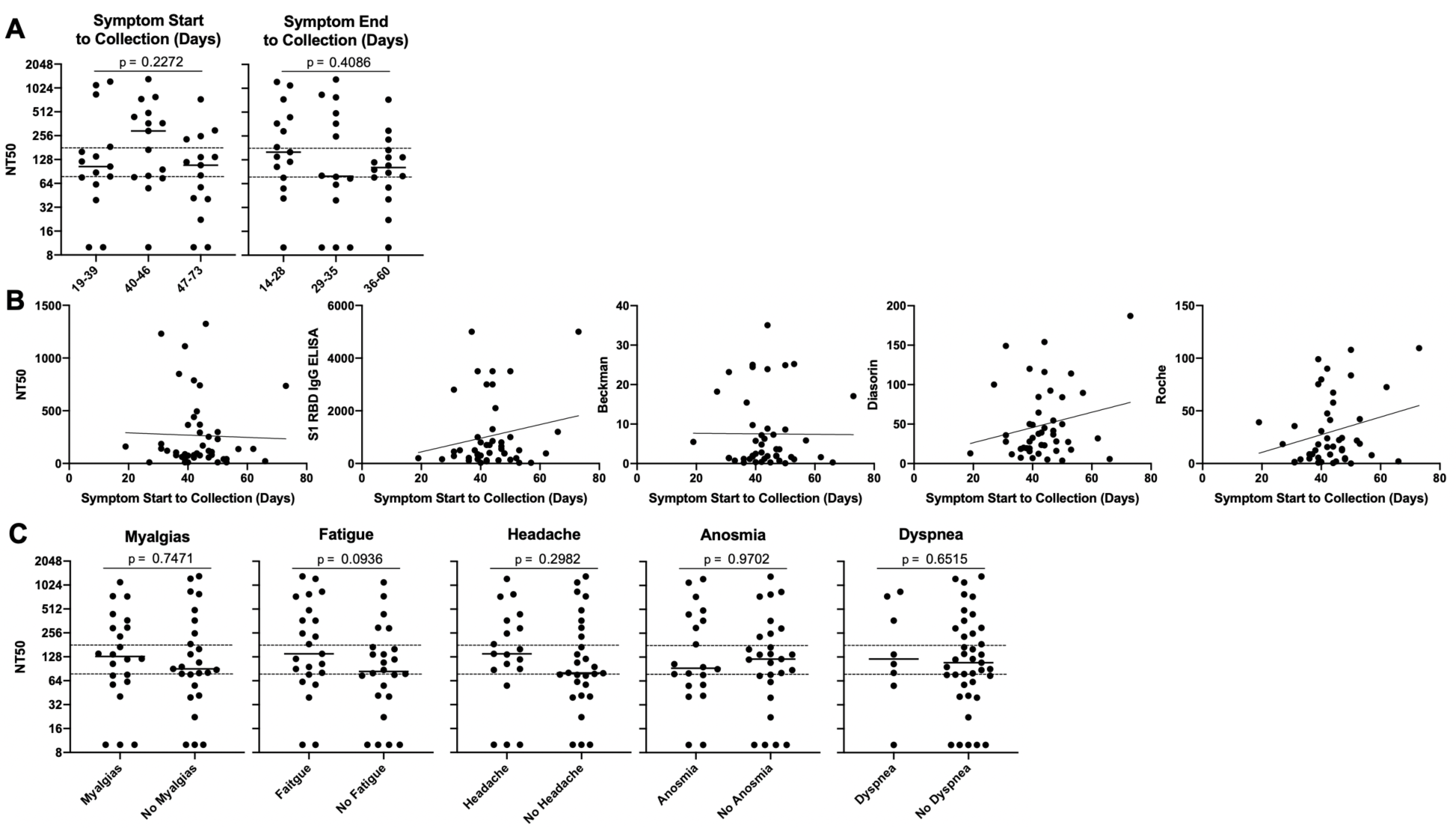
Donor symptomatology and timing of plasma collection. (A) Relationship between neutralization activity and duration of time between date of symptom onset or end, and date of plasma collection, as divided into tertiles. Kruskal Wallis test was used to determine statistical significance. (B) Linear regression analysis of days from symptom onset to plasma collection and neutralization activity. (C) Relationship between presence or absence of five most commonly reported symptoms (fever excluded) and neutralization activity. Mann-Whitney U test was used to determine statistical significance.

## Notes

Funding: This work was supported by UMN Department of Medicine funding (TDB), National Institutes of Health 1R01AI153602 (VDM). The NorthShore COVID-19 convalescent plasma collection program was supported by private donations to the NorthShore Foundation.

### Competing Interest Statement

The authors have declared no competing interest.

